# An Open Dataset of Human Electroencephalographic and Behavioural Responses to Food Images

**DOI:** 10.1101/2025.11.07.687287

**Authors:** Violet J. Chae, Tijl Grootswagers, Stefan Bode, Daniel Feuerriegel

## Abstract

Investigating the neurocognitive mechanisms underlying food choices has the potential to advance our understanding of eating behaviour and inform health-targeted interventions and policy. Large, publicly available neural and behavioural datasets can enable new discoveries and targeted hypothesis tests, yet no such datasets are currently available. We present the FOODEEG dataset containing electroencephalographic (EEG) responses to a diverse array of food images, as well as behavioural measures of food cognition (food categorisation task, food go/no-go task, and food choice task responses), collected from 117 participants. We also provide normative ratings for the food image stimuli with respect to 22 food attributes, including nutritive, hedonic and taste properties, familiarity, and elicited emotions. Our dataset also includes questionnaire-based measures of participants’ food motivations, dietary styles, and general motivational tendencies. In the validation analyses, we demonstrate that early food-evoked EEG responses in our dataset are consistent with observations in previous work. The FOODEEG dataset will be valuable for accelerating research into the neural substrates of visual food processing, dietary decisions, and individual differences.

## Background & Summary

Food-related decisions are highly complex, often involving the integration of sensory and mnemonic information as well as the influence of social, cognitive, and contextual factors. Poor diet has been linked to a number of negative health consequences, including increased risks for cardiovascular and neurodegenerative diseases, and certain types of cancer^1,2^; yet, maintaining a healthy diet is a challenge for many people. While recent neuroimaging work has shed light on the mechanisms underlying food-related cognition, such as the visual processing of foods^3–8^, cognitive reappraisal^9–14^, working memory^15–17^, and reward processing^18–20^, we do not yet fully understand how people make dietary decisions. Gaining insight into these neural processes could inform future interventions aimed at improving dietary choice, and subsequent health and wellbeing.

Using visual food cues (e.g., food images) to investigate the neural mechanisms of dietary choice confers several advantages. Compared to using real foods as stimuli, using food images allows for precise characterisation of stimulus features (e.g., size, colour), increases reproducibility across and within studies, and reduces cost. There is also evidence that responses to food images are associated with eating behaviour. A meta-analysis^21^ reported that cue reactivity and craving elicited by visual food cues predicted eating behaviour and weight gain with a similar effect size as those elicited by exposure to real foods, and with a greater effect size than those elicited by olfactory food cues. Researchers can also readily access food image stimulus sets for use in experimental research, such as Food-pics^22,23^, the FoodCast Research Image Database^24^, the Cross-Cultural Food Image Database^25^, the Open Library of Affective Foods^26,27^, and the Macronutrient Picture System^28^.

These image databases can be used to systematically measure neural responses to foods that vary across a wide range of choice-relevant dimensions. However, collecting a large amount of neuroimaging data using these stimulus sets can be time consuming and expensive. Recently, large-scale neuroimaging datasets collected while viewing scenes, objects, and animals have become publicly available (e.g., Natural Scenes Dataset^29^; THINGS-EEG^30^), allowing for a rapid expansion in research on the visual processing of objects and scenes. Providing large, high-quality datasets containing neural responses to visual food cues can similarly help researchers to devise and test novel questions in food cognition research and nutrition science. Such datasets also enable replication analyses in conjunction with other neuroimaging datasets to test the generalisability of novel findings. Furthermore, large-scale behavioural and neural datasets can be used to train models of eating behaviour to predict health outcomes^31^.

Food choice behaviour can also vary widely, and may reflect individual differences in eating styles^32–34^. Such differences may also be reflected in neural activity. Examination of neural responses to food images could potentially reveal mechanisms that underpin individual differences in dietary choices. Differences in food-evoked neural responses has been reported for people who differ in their levels of restrained eating (the tendency to restrict food intake to control body size or weight)^35–37^, external eating (tendency to eat in response to external food cues)^38,39^, and emotional eating (tendency to eat in response to negative emotions)^40^. Individuals can also vary in their motivations when making food choices, with different levels of consideration given to factors such as health, mood, pleasure, familiarity, habits, and hunger when choosing what to eat^41,42^. However, few studies have examined the associations between food motivations and the neural processing of food cues. Another important facet of food cognition is inhibitory control. A recent meta-analysis^43^ revealed that food-related inhibitory control (as measured using laboratory-based behavioural tasks) plays a role in food consumption, though the strength of the association was small. However, few studies have included measures to examine similar differences in neural responses^39^. Large datasets are particularly important when examining the links between neural processing of foods and individual differences in eating style, food motivations, and inhibitory control abilities, given the high power required for between-subject group comparisons or correlational analyses.

Here, we present a large dataset (*N* = 117) comprising electroencephalographic (EEG) and behavioural responses to a broad set of 120 food image stimuli. This dataset includes EEG data and behavioural responses recorded during a food categorisation task, and also behavioural data collected during a food go/no-go task and a paired food choice task. Questionnaire-based measures of participants’ food motivations (Food Choice Questionnaire^42^, Eating Motivation Survey^41^), dietary styles (Dutch Eating Behaviour Questionnaire^34^, Three Factor Eating Questionnaire-R21^33,44^), general motivational tendencies (Behavioural Activation and Behavioural Inhibition Scales^45^), and hedonism (Present Time Orientation Scale^46^) are included. Additionally, we provide normative ratings on 22 food attributes for each of the food images, spanning nutritive properties (healthiness, calorie content, edibility, and level of transformation), hedonic properties (tastiness, willingness to eat, positive and negative valence, and arousal), familiarity (previous exposure, recognisability, and typicality), taste properties (sweetness, saltiness, sourness, bitterness, and savouriness), and emotional properties (happiness, surprise, disgust, craving, and guilt) collected from separate online samples (total *N* = 624).

Users may take advantage of the high temporal resolution afforded by EEG, which allows for the characterisation of rapid neural processes that underlie visual perception and decision-making. Previous work has shown that abstract information about objects beyond low-level visual features, such as animacy, naturalness, and elicited emotions, are reflected in EEG signals within the first several hundred milliseconds after image onset^47–49^. This dataset is well suited for uncovering the precise timing of neural processes that occur in response to food images.

The EEG task was carefully designed to separately measure the appraisal and decision stages of food evaluation (Figure 1A). In this task, participants were presented with a food image and were asked to evaluate the food on one of three attributes: healthiness, tastiness, and willingness to eat. Each food image was presented three times (once for each attribute). We sought to separate the appraisal and decision stages of food evaluation by including a viewing phase, in which participants viewed the food image in isolation for 2 seconds, before the response phase, in which participants were presented with the prompt indicating the attribute they were to evaluate the food on. The mapping of the response options onto response keys was randomised across trials to prevent anticipatory motor preparation. We reduced priming effects by presenting healthiness, tastiness, and willingness to eat trials in a randomised order within blocks, rather than presenting each trial type in separate blocks. This ensured that the participants were not able to anticipate which of the three attributes would be judged during food image viewing. This task design allows for investigations into the neural mechanisms underlying food appraisal and the visual processing of foods while minimising the influences of neural signals related to decision or motor preparation. Users interested in the neural correlates of food-related decision-making processes can instead examine relationships between choice behaviour (e.g., response times) and neural activity in the response phase. We recommend conducting analyses on the attribute (e.g., healthiness, tastiness) or category (e.g., meals, snack foods) level, rather than at the level of individual food images, due to the low number of trials per image (3 trials).

**Figure 1.**
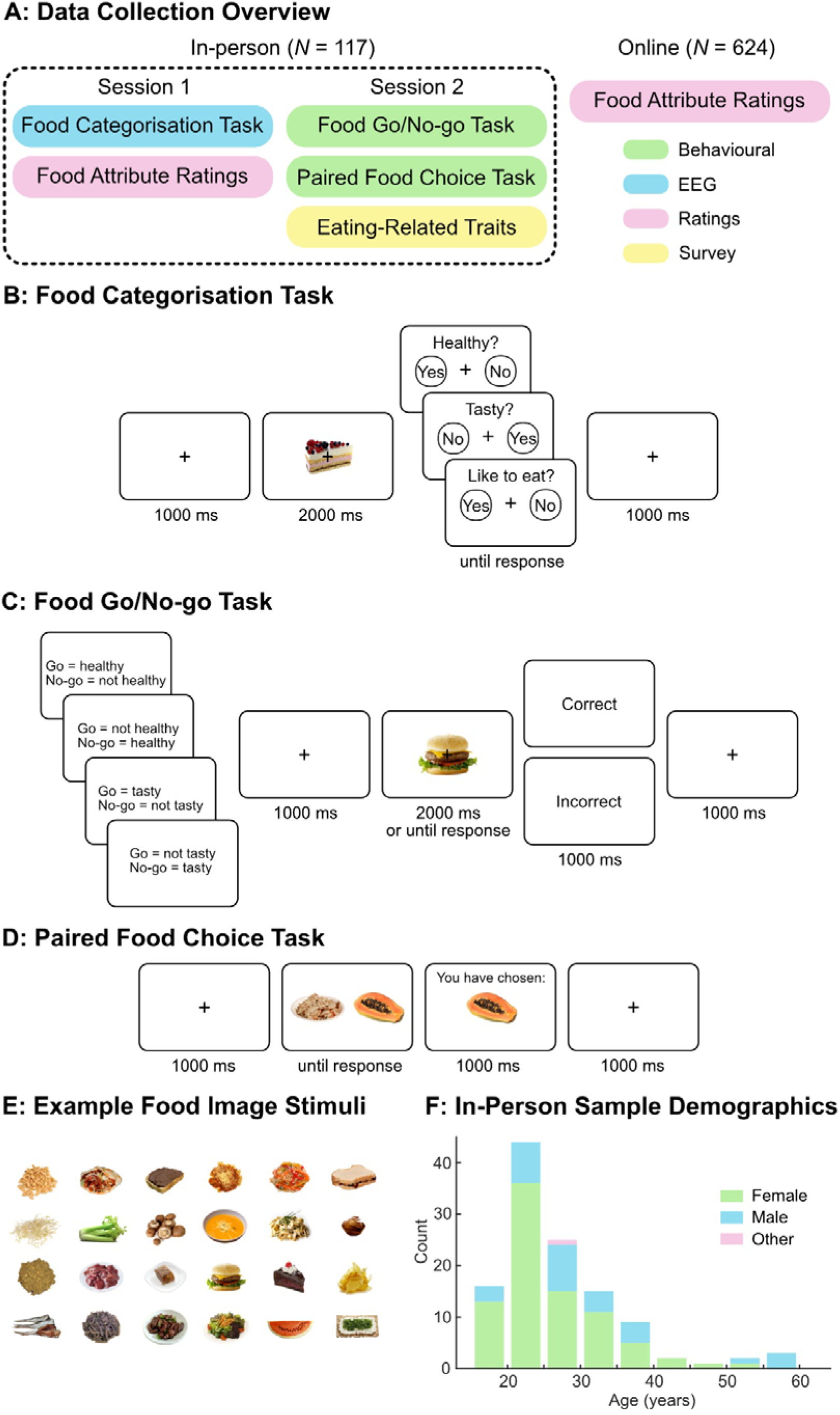
Overview of data collection and experimental trial diagrams. A) Data collection overview. The data was collected over two in-person sessions and online. In the first in-person session, participants completed the food categorisation task (while EEG was recorded) and provided ratings on the food images for three attributes. In the second in-person session, participants completed two behavioural tasks (food go/no-go task and paired food choice task) and filled out survey measures of dietary styles, food motivations, and general motivational tendencies. In an online experiment using a separate sample, attribute ratings were collected for the food images for 22 choice-relevant food attributes. B) Food categorisation task. For each trial, participants viewed a food image (2 s), followed by a response screen with one of three prompts probing healthiness, tastiness, or willingness to eat and one of two keyboard press response mappings (‘z’ key for Yes and ‘m’ key for No, or the reverse). Participants were instructed to respond as quickly as possible. C) Food go/no-go task. In each trial, participants were presented with a food image and either responded as quickly as possible with a keyboard press for Go foods or withheld any responses (2 s) for the No-go foods. Feedback (‘Correct’ or ‘Incorrect’) was given after each trial. Participants completed four blocks with different Go and No-go foods. D) Paired food choice task. Participants were presented with two food images and were asked to choose the food that they would prefer to eat more, followed by a feedback screen showing the food image that they had selected. E) Example food image stimuli. Food images were selected to include a wide variety of food types and also included foods generally thought to be less appetising or familiar. F) In-person sample demographics. Age distribution for the in-person sample (*N* = 117) is shown in a histogram. Green, blue, and pink denote the proportions of participants who identified as female, male, and other respectively.

The food image stimuli were carefully curated to ensure that there was substantial variance in key food attributes. This is often difficult to achieve because people tend to view most foods as appetising and positively valenced. Ninety-one images were selected from an established food image database (Food-pics^23^) and consisted of mostly palatable and familiar foods. Twenty-nine images of foods that are less palatable or familiar, but still edible, were sourced from online searches and included in the stimulus set. Some existing work has compared behavioural responses towards edible versus inedible foods (e.g., rotten or decayed foods^50,51^), but few studies have included foods that are unappetising but not harmful to eat in their stimuli.

Rating data on a wide range of food attributes indicate that our stimuli sufficiently varied on key attributes that are relevant for dietary choice. This indicates that our dataset is suitable for analyses that are sensitive to fine-grained differences in the variables of interest, such as multivariate pattern analysis (MVPA) or representational similarity analysis (RSA). For example, MVPA can be applied to investigate how fine-grained rating information about a food attribute of interest covaries with patterns of EEG data, to delineate the precise neural time-courses of these representations during food viewing (see Technical Validation for an example). Alternatively, fine-grained rating information can be used to construct models of similarity for a given food attribute and compare them to models of similarity for neural or behavioural data. Recent work has demonstrated how RSA can be applied to this dataset.^3^

Beyond what we provide here, additional ratings can be easily collected from new online participant samples to assess an even wider range of food attributes and how they covary with neural responses in our dataset. Users may also use these normative ratings in combination with the food images to inform stimuli selection, to ensure a particular distribution of attributes to suit their research goals.

In sum, the FOODEEG dataset comprises of a rich array of neural and behavioural measures during food cognition tasks collected from a large sample (*N* = 117). Given the multifaceted nature of eating behaviour, we have included measures of eating styles, food motivation, and relevant personality traits which may be useful in controlling for, or investigating, individual differences. We also provide a comprehensive set of ratings on 22 food attributes that encompass nutritive, hedonic, familiarity, taste, and emotional properties of foods. The FOODEEG dataset has the potential to advance food cognition research by supporting a wide range of research projects, including investigations linking neurocognitive mechanisms to individual differences, food attribute processing, and the dynamics of choice behaviour.

## Methods

Neural and behavioural data were collected in-person over two testing sessions. In Session 1, participants performed a food categorisation task (Figure 1B) in a testing booth while EEG was recorded. Participants then rated the same food images on healthiness, tastiness, and willingness to eat. In Session 2, participants completed a food go/no-go task (Figure 1C) and a paired food choice task (Figure 1D). They also completed questionnaires relating to their eating styles, food-specific motivations, and general motivational tendencies. A separate sample of participants completed an online food rating task, in which participants were asked to rate the food image stimuli on multiple food attributes. Figure 1A shows the overview of the data collection procedure. This study was approved by the Human Research Ethics Committee of the University of Melbourne (HREC ID 24850).

### Participants

For the in-person sessions, participants (*N* = 117) were recruited through online study advertisements on the University of Melbourne student portal and via posters placed around the University of Melbourne Parkville campus. The advertisements contained a link to an online survey regarding the eligibility criteria and those that met the requirements were contacted through email. Participants were eligible to participate if they were at least 18 years old, were fluent in written and spoken English, did not have a history of any eating disorders, were not on a calorie restriction diet, and did not have any dietary restrictions (e.g., for health or religious reasons). Participants were instructed to refrain from eating any food for a minimum of three hours prior to both in-person sessions. Participants were reimbursed AUD $50 after completing both sessions. The sample had a mean age of 26.8 years (SD = 8.50, range 18-57 years). 84 participants identified as female, 32 as male, and one as other. Note that a large proportion of our in-person sample consisted of female participants. As gender and sex-related differences have been reported in some aspects of dietary behaviour (e.g., eating styles and food craving)^52^, researchers who wish to examine such processes may consider taking a balanced sample from the full dataset or opt for using data from the female participants only.

Online participants (*N* = 624) were separately recruited through the University of Melbourne Research Experience Program, which offers research participation opportunity to undergraduate students in the Melbourne School of Psychological Sciences. Participants took part in the study to earn course credit. All online participants were at least 18 years old and fluent in written and spoken English. Participants indicated that they did not have a history of any eating disorders, were not on a calorie restriction diet, did not have any dietary restrictions, and had not previously taken part in the in-person testing sessions. The online sample had a mean age of 19.55 years (SD = 2.92, range 18-53 years). 482 participants identified as female, 132 as male, 7 as non-binary/other, and 3 preferred not to disclose. Course credit was allocated to those who participated in the online experiment in full.

All participants were given a Plain Language Statement detailing the study procedure, potential risks involved, and how their data would be stored and shared. In-person participants were informed of the possibility of minor discomfort from wearing the EEG cap and fatigued during the task, while online participants were informed that there were low to negligible risks. All participants were advised that their information would be kept confidential subject to legal requirements and that the data from the study may be used in future research and may be made publicly accessible in a de-identified format. Participants signed a consent form after reading the Plain Language Statement. Prior to the deposition of the data in the repositories, all data were deidentified to protect participant anonymity. Specifically, each participant’s name and contact information was removed and replaced by an arbitrary ID number.

### Stimuli

The stimuli consisted of 120 colour images of food items against a white background. Ninety-one images were drawn from the Food-pics database^23^ and 29 images were drawn from online search. Food-pics^23^ is a database of food images containing images of a wide variety of food items designed for use in research on eating behaviour. The 91 food images were selected to include a wide range of food types (e.g., vegetables, fruits, savoury dishes, sweet dishes, meats). Food images that portrayed multiple types of foods or included cutlery were not included. An additional 29 images were obtained from image hosting websites (e.g., Flickr, Adobe Stock). We aimed to increase variance in key food attributes (e.g., tastiness, valence, familiarity) in our stimuli set. To this end, we included images of foods that were generally less palatable but were nevertheless edible. Examples of the food images are displayed in Figure 1E.

We note that foods naturally vary on low-level visual features (e.g., colour, contrast, and visual complexity) which may covary with other aspects of foods, such as healthiness, level of transformation, or valence. For example, vegetables make up a large proportion of foods that are green and at the same time, tend to be judged as healthy. Processed foods tend to be more visually symmetrical compared to raw foods. Such low-level image properties are known to reliably modulate early neural responses (e.g., N1, P1)^53,54^. However, fully controlling for these image properties is difficult to achieve while keeping the stimuli recognisable and ecologically valid. We encourage users to control for relevant low-level visual features in their analyses^3^. To support this, we have included luminance and contrast values for each food image in the Stimuli directory on the Open Science Framework repository^55^. Luminance and root mean square contrast values were calculated as the mean and standard deviation of the greyscale values of the image, respectively^56^. These image properties are provided to help users with stimuli selection or for inclusion into analyses to control for neural activity evoked by these low-level visual features. Users can extract other image properties using the full set of food images provided in the Stimuli directory on the Open Science Framework repository^55^.

### Procedure

Participants attended the two in-person sessions on different days, approximately 1 week apart. In Session 1, participants carried out a food categorisation task presented using Psychtoolbox^57^ interfacing Matlab R2022a. Next, participants evaluated each food image on three food attributes (healthiness, tastiness and willingness to eat) using Qualtrics. During the second session, participants completed two behavioural tasks: a food go/no-go task and a paired food choice task presented using PsychoPy^58^ interfacing Python (v3.11). All experimental code are available via the Open Science Framework^55^. Participants also completed questionnaires relating to dietary styles, food motivations, and general motivational tendencies using Qualtrics.

#### Food categorisation task with EEG recording

During the food categorisation task, each food item was presented to the participants three times (once per trial type: healthiness, tastiness, and willingness to eat) while EEG data was collected (Figure 1A). Food images were presented centrally on an ASUS ROG Swift PG258Q monitor (24.5 inch, 1920 × 1080 pixels, 60 Hz refresh rate), subtending 6.6 × 3.5 degrees of visual angle. To minimise head movement, participants were seated approximately 50 cm from the screen with their head stabilised using a chin and forehead rest. Each trial started with the presentation of a fixation cross for 1 s, after which the food image appeared alongside the fixation cross for 2 s. The participants were then shown a question prompt corresponding to the trial type: “Healthy?” in a healthiness trial, “Tasty?” in a tastiness trial, and “Like to eat?” in a willingness to eat trial. Response mapping was presented alongside the prompt to indicate which of the left (‘z’) and the right (‘m’) keyboard keys represented a ‘Yes’ and ‘No’ responses. The assignment of keys to responses was equally likely to be the left keyboard press indicating a ‘Yes’ response and the right keyboard press indicating a ‘No’ response, as the reverse left/right mapping, on any given trial. For each participant, the trial types (healthiness, tastiness, and willingness to eat) were intermixed and presented in a randomised order, as was the order of food images within the sequence. The randomisation of the trial type order and response mapping prevented participants from planning a keypress motor action in advance during the image presentation period. Participants were instructed to respond as quickly and accurately as possible, and to monitor the switching of the response mapping carefully as it could change for each trial. Prior to the main experiment, participants completed five practice trials and confirmed that they had understood the task. To minimise eye movement artifacts, participants were asked maintain fixation on the central cross, which remained on screen throughout each experimental block. Response time (the interval between the prompt and response mapping presentation and the participant’s keypress) was recorded for each trial. Participants completed six blocks of 60 trials each, with self-paced breaks between each block. The task took approximately 45 minutes to complete, including breaks.

EEG was recorded using a Biosemi Active II system (Biosemi, The Netherlands) with 64 scalp electrodes at a sampling rate of 512 Hz using common mode sense and driven right leg electrodes (http://www.biosemi.com/faq/cms&drl.htm). Eight additional electrodes (two electrodes placed 1 cm from the outer canthi of each eye, four electrodes placed above and below the centre of each eye, and two electrodes placed above the left and right mastoids) were also attached. Throughout the recording, electrode offsets were maintained within the range of ±20 *µ*V.

#### Healthiness, tastiness, and willingness to eat ratings

After the food categorisation task, participants rated each food image on healthiness (“How healthy is this food?”), tastiness (“How tasty is this food?”) and how willing they were to eat it (“How much would you like to eat this food?”). Participants were asked to use a computer mouse to indicate their response on a continuous rating scale ranging from “Not at all” to “Very much”. We randomised image presentation order for each participant, while ensuring that the same food image was not presented across two consecutive trials.

#### Food go/no-go task

Adapted from the standard go/no-go paradigm, the food-specific go/no-go task has been used in previous research to measure food-related inhibitory abilities^59,60^. Stimulus set selection was carried out for each individual participant, using their attribute ratings from Session 1. Healthiness and tastiness ratings from Session 1 were used to select 60 food image stimuli across four categories: 15 Most-Healthy foods, 15 Least-Healthy foods, 15 Most-Tasty foods, and 15 Least-Tasty foods. To select the Most-Tasty foods, 15 foods with the highest tastiness ratings were selected. To select Least-Tasty foods, 15 foods with the lowest tastiness ratings were selected. From the remaining foods, Most-Healthy and Least-Healthy foods were selected in the same way. If there were more than 15 foods with the highest or lowest ratings for an attribute, 15 were randomly selected from those foods. No foods were included in multiple categories. Prior to the start of the task, participants were shown all 60 food images and the category each belonged to. It was indicated to the participants that the foods were categorised according to their own ratings from the previous session, and they were asked to let the experimenter know if they disagreed with the categorisations for any of the foods. If the participant disagreed with the categorisation of a food, the food was removed from the category and replaced with the food with the highest (for Most-Healthy and Most-Tasty categories) or the lowest (for Least-Healthy and Least-Tasty) rating on the appropriate attribute out of the unselected foods.

Participants completed four blocks of the go/no-go task. In the first block, Most-Healthy foods served as the Go stimuli and the Least-Healthy foods served as the No-go stimuli. In the second block, Most-Tasty foods served as the Go stimuli and the Least-Tasty foods served as the No-go stimuli. The Go and No-go stimuli in the first and second blocks were switched for the third and fourth blocks respectively. There were 100 trials in each block. In each block, Go foods were presented in 75% of the trials and No-go foods were presented in 25% of the trials. At the start of each block, instruction text indicated to the participants which food categories served as the Go and No-go stimuli for the block. Next, the participants were shown a series of food images and were asked to make a response via keyboard press as quickly as possible to the Go foods and to withhold the response to the No-go foods. A fixation cross was presented for 1 s between trials. Block order was randomised for each participant.

#### Paired food choice task

To assess people’s preferences across foods that differed in attributes (e.g., healthiness and tastiness), participants selected which food they would prefer to eat more out of pairs of foods they had previously indicated they would be willing to eat. Stimuli selection was carried out for each individual participant, using their attribute ratings from Session 1. Out of the food images that had been rated over 50 (scale: 0-100) on willingness to eat in Session 1, 25 food images were selected. If less than 25 food images had been rated over 50 on willingness to eat, the subsequent highest rated food images were included in the stimuli to bring the total number of food images to 25.

Participants completed five blocks of the paired food choice task. There were 60 trials in each block. In each trial, participants were presented with two food images on the left and right side of the screen. Participants indicated which food they would prefer to eat more with a keyboard press (‘z’ to select the left food and ‘m’ to select the right food). Every possible pairwise combination of the 25 food images was presented once, totalling 300 trials. A fixation cross was presented for 1 s between each trial.

#### Questionnaires

Participants’ dietary styles, food motivations, general motivational tendencies, and hedonism were measured using self-report scales. Dietary styles were assessed using the Dutch Eating Behaviour Questionnaire^34^ and the Three Factor Eating Questionnaire Revised 21-Item^33,44^. Food-related motivations were assessed using the relevant subscales of the Food Choice Questionnaire (Health, Mood, Sensory Appeal, Natural Content, Weight Control, and Familiarity subscales)^42^ and the Eating Motivation Survey Brief (Liking, Habits, Need and Hunger, Health, Pleasure, Natural Concerns, Weight Control, Affect Regulation subscales)^41^. Motivational tendencies were measured using the Behavioural Activation and Behavioural Inhibition Scales^45^ and hedonism was assessed using the Present Time Orientation Scale^46^. The descriptive statistics for the survey measures are presented in Table 1.

**Table 1.**
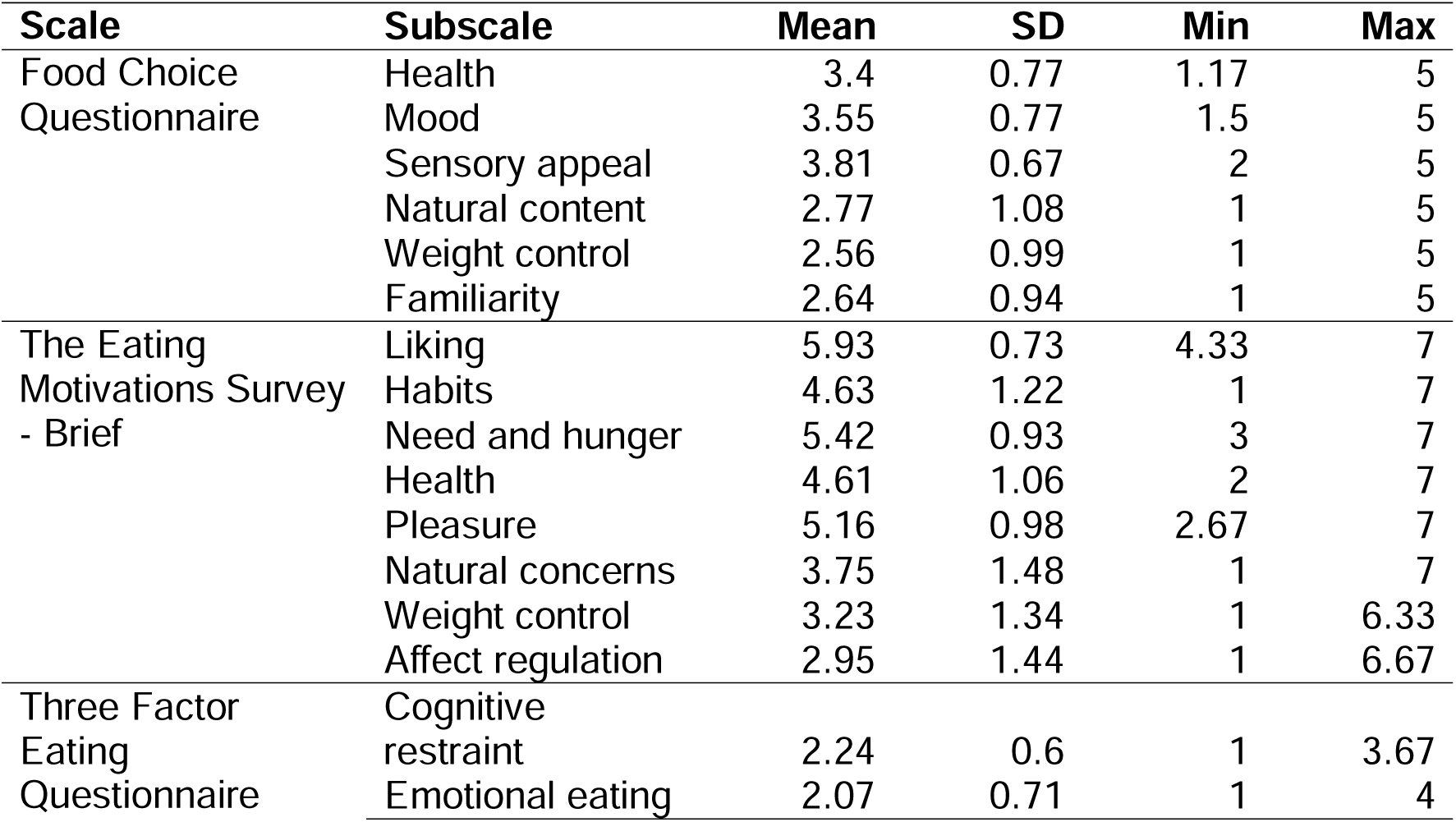
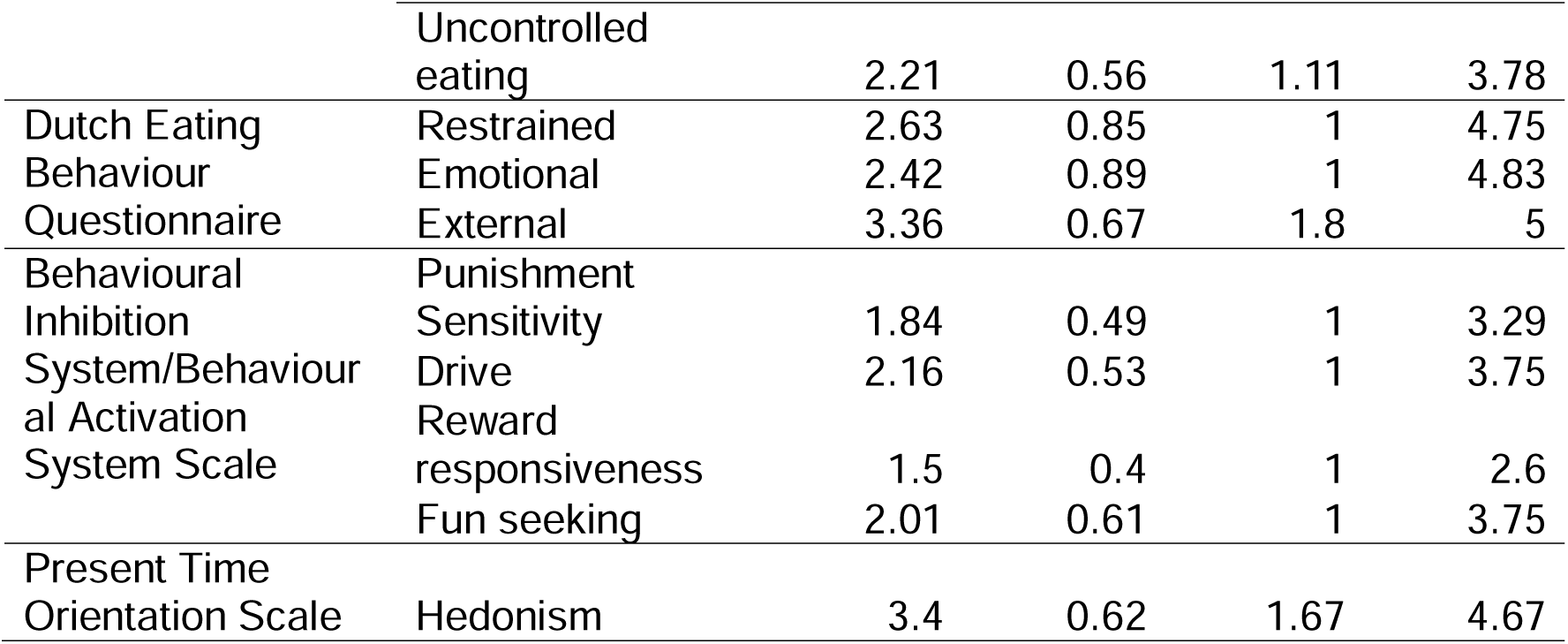
Summary statistics for survey measures of dietary style, food motivations, general motivational tendencies, and hedonism. Mean, standard deviation, and range of individual participant’s mean scores for the subscales of Food Choice Questionnaire, the Eating Motivations Survey – Brief, Three Factor Eating Questionnaire, Dutch Eating Behaviour Questionnaire, Behavioural Inhibition System/Behavioural Activation System Scale, and Present Time Orientation Scale are reported.

#### Online food attribute ratings

A separate sample of participants completed the online rating task. Participants recorded their current hunger (“How hungry are you currently?”) and satiety (“How full are you currently?”) level on a sliding scale from ‘Not at all’ to ‘Very much’ and indicated how much time has elapsed since they had consumed food (“How long ago did you last eat food?”) from one of the following options: ‘Less than an hour ago’, ‘1-3 hours ago’, ‘3+ hours ago’, and ‘I do not know’. Each participant rated a third of the full set of food images (40 out of the 120 images) on 12 (out of 22) food attributes. Each individual food image was evaluated by 204-212 participants on healthiness, tastiness, and willingness to eat. Each image was rated by 134-145 participants on calorie content, level of transformation, edibility, positive and negative valence, arousal, previous exposure, recognisability, and typicality, and by 66-70 participants on emotional properties (happiness, disgust, surprise, craving, and guilt) and taste properties (sweetness, saltiness, bitterness, sourness, and savouriness). Individual participants’ responses and summary statistics for each attribute and each food image (mean, SD, and range) are provided in the Ratings directory on the Open Science Framework repository^55^. For each participant, both the sequence in which food images appeared and the order in which each image’s attributes were evaluated were randomised. Table 2 displays the full list of food attributes and the corresponding prompts and additional information presented to the participants. Participants indicated their rating for each attribute on a sliding scale (0-100) using a computer mouse.

**Table 2.**
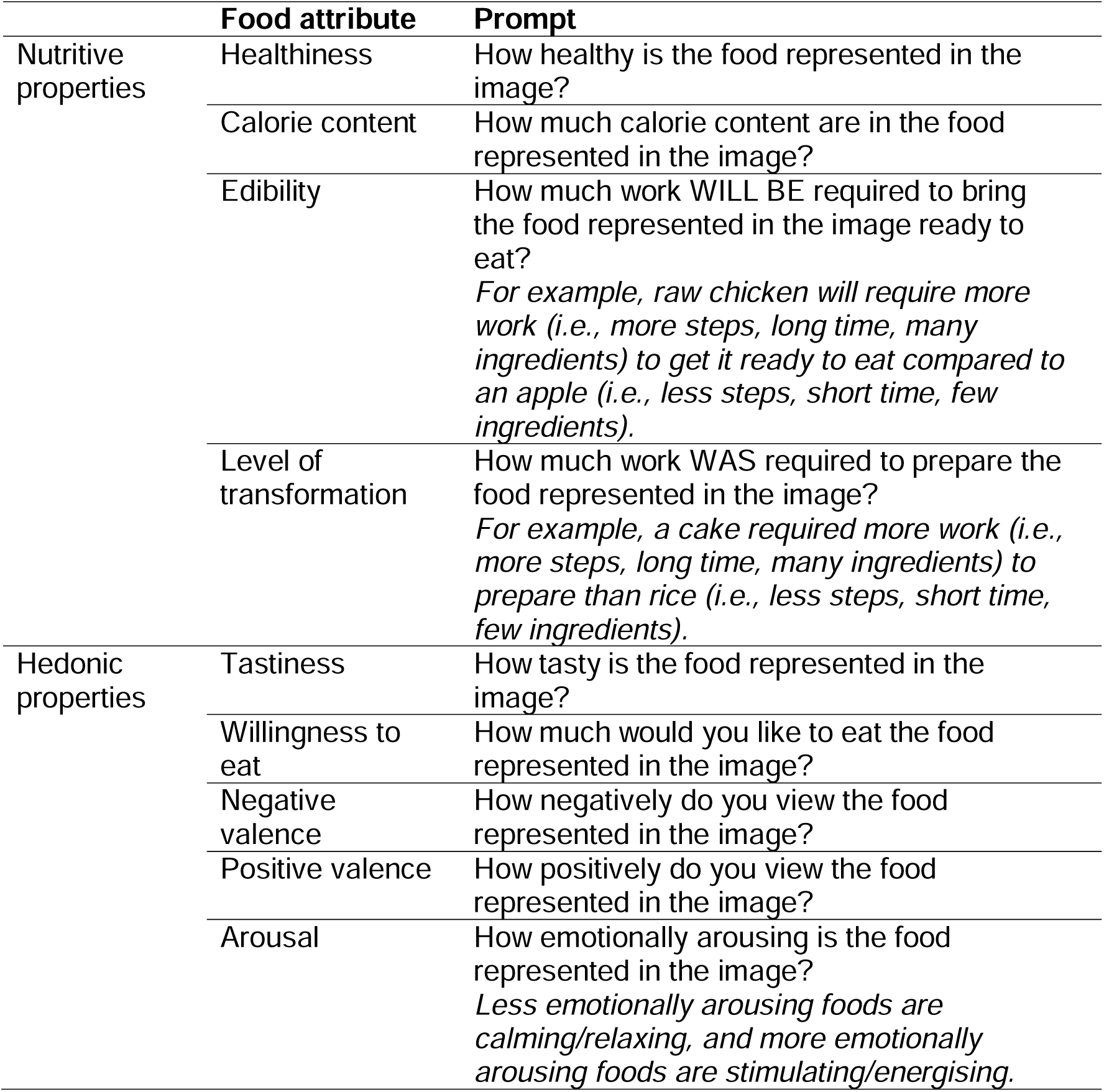
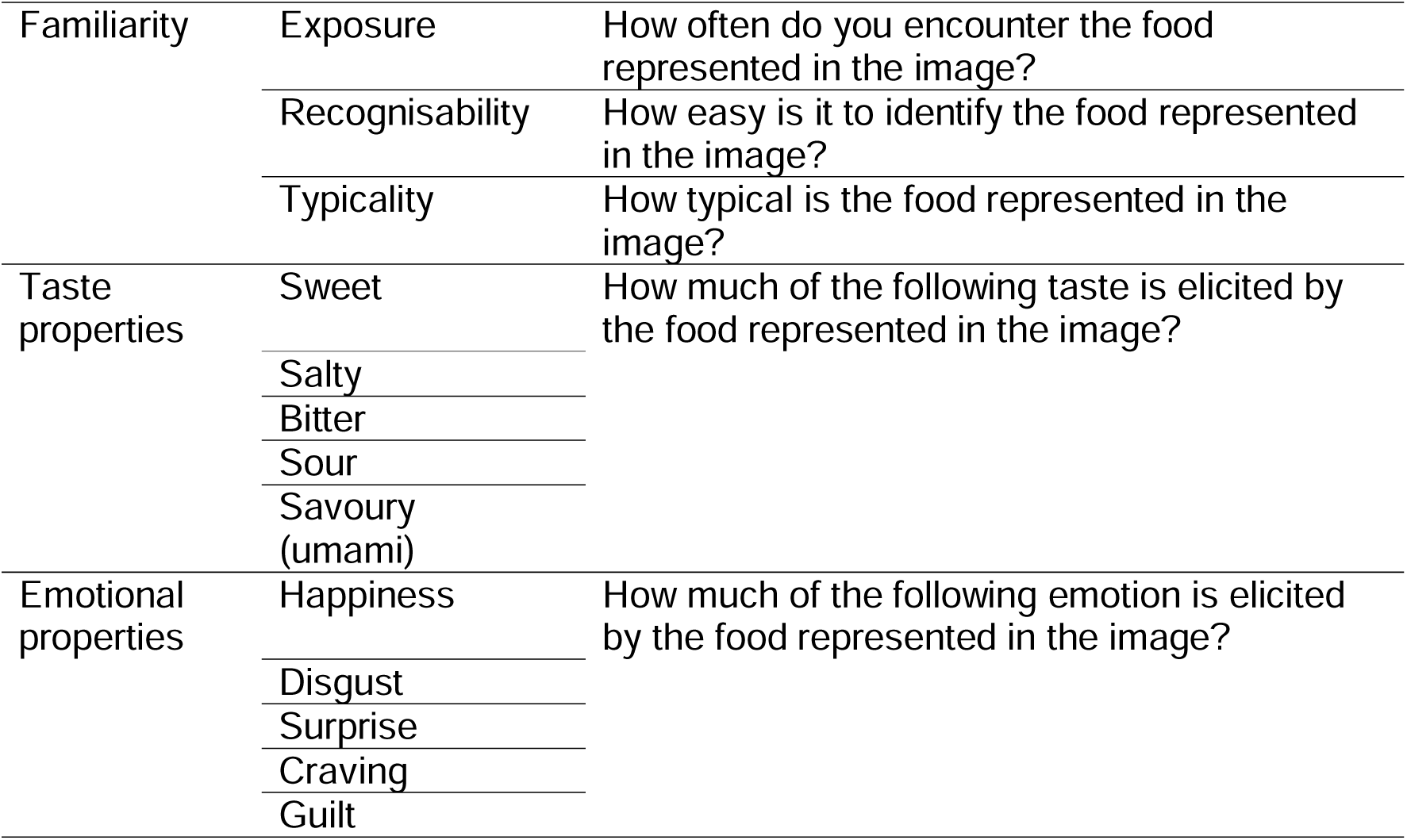
Food attribute ratings and corresponding prompts. Ratings on 22 food attributes across five categories (nutritive, hedonic, familiarity, taste, and emotional properties) were collected for the food image stimuli using the corresponding prompt for each food image. For edibility, level of transformation, and arousal, additional information was displayed alongside the prompt (in italics).

## Data Records

All data and code are publicly available. The raw and processed EEG recordings and behavioural data are hosted in Brain Imaging Data Structure (BIDS) standard^61,62^ on OpenNeuro^63^ (https://doi.org/10.18112/openneuro.ds007012.v1.1.0). The raw EEG recordings are provided in BioSemi data format (.bdf) and the cleaned continuous EEG files are provided in EEGLAB data format (.set). Custom code, food image stimuli, survey data, and normative ratings are available from the Open Science Framework^55^ (https://doi.org/10.17605/OSF.IO/Y9PMF).

At the root level of the OpenNeuro repository^63^, several sidecar files are provided to document metadata that apply to all of the lower levels of the repository. The participant-related TSV file records demographic and descriptive information such as participant ID, age, gender, whether they are suggested to be excluded from analyses, and any additional notes. The accompanying JSON sidecar file provides column-level metadata describing the meaning and units in the TSV file. A dataset description JSON file provides top-level information about the entire dataset, including its name, the BIDS version it conforms to, author information, and links to associated publications or data repositories. An EEG JSON file contains metadata for the EEG recording, including task name, the number of EEG channels, the amplifier and electrode system used, the type of EEG reference and ground electrode, hardware and software filter settings, the recording institution, and the recording type. The events JSON files serve as data dictionaries for the accompanying TSV files containing the raw event-related data. Within each subject folder, data are organised by modality into datatype subdirectories. EEG recordings are stored under eeg/, while the behavioural data from the food go/no-go and paired food choice tasks are stored under beh/. Each subject’s EEG directory contains the raw EEG data file alongside TSV files containing details of all experimental events that occurred during the recording. The derivatives folder contains the cleaned continuous recordings for all subjects.

The Open Science Framework repository^55^ is organised into five top-level directories: Derivatives, Experiment, Questionnaires, Ratings, Stimuli, and Validation. The Derivatives directory stores the code used to process the raw EEG data into cleaned continuous data. The Experiment directory contains the code used to run the three experimental tasks, organised into three subdirectories (food categorisation task, food go/no-go task, and paired food choice task). Self-report questionnaire materials, individual participant responses, and summary statistics are provided in the Questionnaires directory. The Ratings directory contains the food attribute ratings for the 120 food image stimuli, organised into two subdirectories. The in-person subdirectory stores ratings collected from the in-person participants at the end of Session 1, while the online subdirectory stores the ratings collected from the online sample. Demographic information for online participants, survey instruments, and summary statistics are provided in the online subdirectory. The Stimuli directory contains the 120 food images used across all experimental tasks. Of these, 91 were drawn from the Food-pics database^23^ available for non-commercial research use under a CC BY-NC-SA licence, and 29 were sourced from online sources. The code used in the technical validation analyses and copies of the figures are included in the Validation directory. See the readme file in the Open Science Framework repository for further details.

## Technical Validation

### EEG data validation

For technical validation of the current dataset, we used two complementary analyses to demonstrate that both binary categorisations and continuous ratings on food attributes covary with patterns of EEG data. We examined EEG responses following food image onset to assess whether event-related potential (ERP) waveforms differed across different food items that categorically varied in attributes (e.g., Healthy versus Not-Healthy foods). Furthermore, we used MVPA to demonstrate that fine-grained information about food attributes is also present in the EEG signals.

#### Data processing

We processed the EEG data using EEGLab v2022.1^64^ interfacing Matlab (R2022a). Channels with excessive noise were identified via visual inspection (mean number of bad channels = 0.26, range 0–5) and omitted from average reference calculations and independent components analysis (ICA). Data segments containing excessive noise were also identified through visual inspection and removed. Five participants were excluded from the analyses due to noisy data or missing trials. Two additional participants were excluded due to high incongruence between their responses in the food categorisation task and their subsequent continuous food attribute ratings. The participant-related TSV file in the OpenNeuro repository indicates which participants are suggested to be excluded. The data were referenced to the average of all channels, excluding those that were excessively noisy. We removed one channel (AFz) to address the rank deficiency introduced by average referencing. A 30 Hz low-pass filter was applied to the EEG data (EEGLab Basic Finite Impulse Response Filter New, default settings). Next, a copy of the dataset was created for the purpose of ICA, to which a 0.1 Hz high-pass filter (EEGLab Basic Finite Impulse Response Filter New, default settings) was applied to improve stationarity. ICA was performed on this copied dataset using the RunICA extended algorithm^65^. The resulting independent component information was subsequently copied to the original dataset^66^. Independent components associated with eye movement artifacts (blinks and saccades) were identified and removed in accordance with established protocols^67^.We interpolated the excessively noisy channels and AFz using spherical spline interpolation. EEG data were segmented into epochs spanning −100 ms to 1000 ms relative to food image onset, and each epoch was baseline-corrected using the data from the 100 ms time window preceding food image presentation. Epochs from trials in which the participant’s response time exceeded 5 s, or in which the amplitude at any channel exceeded ±150 *µ*V, were excluded from further analyses. On average, 346 out of 360 trials were retained per participant (range 283 – 360).

#### Event-related potentials

We demonstrate that ERP waveforms closely resemble those observed in previous work using visual food stimuli^6,8,11,68–70^, and that differences in ERPs can be observed across food images that vary categorically in attributes of interest (e.g., Healthy versus Not-Healthy, Tasty versus Not-Tasty foods). Figure 2 shows group-average ERPs measured at occipital electrodes (Oz, O2, O1, and Iz) and at parietal electrodes (Pz, P1, P2, CPz, and POz) following food image onset. For each participant, we split the trials according to the participants’ responses during the food categorisation task on healthiness and tastiness trials (Healthy versus Not-Healthy, Tasty versus Not-Tasty). We then averaged EEG responses across trials to derive ERPs for each choice condition. ERPs were averaged over electrodes within each region of interest (occipital and parietal) to further improve the signal-to-noise ratio. We compared ERPs following Tasty and Not-Tasty food images, and Healthy and Not-Healthy food images, using cluster-based permutation tests to correct for multiple comparisons (10,000 permutation samples, cluster forming alpha = .01, family-wise alpha = .05) using functions from the Decision Decoding Toolbox v1.1.5^72^.

**Figure 2.**
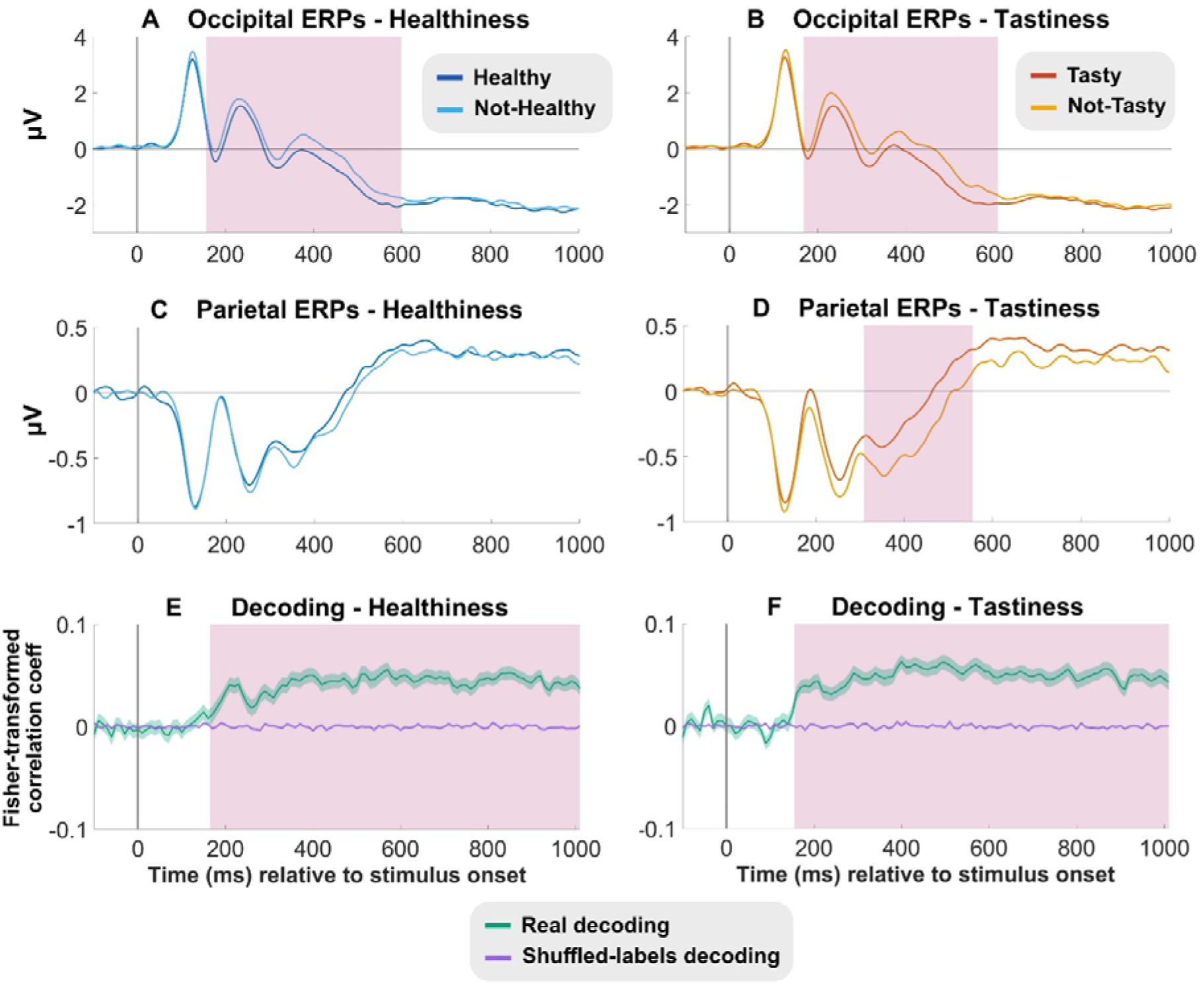
Group-average ERPs and decoding results following food image onset. The top row shows ERPs evoked at occipital (Oz, O2, O1, and Iz) electrodes by A) Healthy (dark blue) and Not-Healthy (light blue) foods and by B) Tasty (red) and Not-Tasty (yellow) foods. The second row displays ERPs evoked at parietal (Pz, P1, P2, CPz, and POz) electrodes by C) Healthy and Not-Healthy foods and by D) Tasty and Not-Tasty foods. For the ERPs, pink shaded areas indicate the time windows at which the mean amplitudes between conditions significantly differed after applying multiple comparisons corrections. The third row shows the decoding results. Using support vector regressions, E) healthiness and F) tastiness ratings were predicted from the EEG data from the time relative to food image onset. Green lines indicate the real decoding results (Fisher-transformed correlation coefficients between predicted and true attribute ratings) and purple lines indicate the shuffled-labels decoding results (empirical chance distribution). Shaded areas around the green and purple lines indicate the standard error of the mean. For the decoding results, pink shaded areas indicate the time windows at which the real decoding results significantly differed from the shuffled-labels decoding results and after applying multiple comparisons corrections.

We observed typical patterns of visual evoked ERPs at occipital channels, including the visual P1, N1 and P2 components that occur within the first 250 ms following food image presentation^11,68,69^. We also observed a later, sustained positivity at parietal channels, resembling the P3 or Late Positive Potential that has been measured in prior work^6,68,69,73^. At occipital electrodes, we observed more negative-going ERP waveforms both for tasty and healthy foods between approximately 150-600 ms (Figure 2A, 2B). At parietal channels, amplitudes were more positive-going between approximately 300-550 ms following Tasty compared to Not-Tasty foods (Figure 2D) as also reported in prior work^6,68,69^. This demonstrates that EEG responses evoked by food images do broadly differ across food attribute decision outcomes, and that it could be used to test more targeted hypotheses relating to food-evoked ERPs.

#### Multivariate pattern analysis

We demonstrate using MVPA that fine-grained food attribute information beyond binary categorisations is also decodable from the current dataset. Multivariate analysis methods (e.g., MVPA, representational similarity analysis) are becoming more widely used in neuroimaging studies, due to high sensitivity that allows for the detection of distributed patterns of neural activity. Previous work has shown that such methods are useful for investigating the fine-grained neural time-courses of food attribute processing^3–5^. The MVPA results suggest that there are patterns in the EEG data that covaried with continuous food attribute ratings and reveal that the current dataset is suitable for multivariate analysis techniques.

Linear support vector regression (SVR) models were implemented using the Decision Decoding Toolbox v1.0.5^72^ to examine whether continuous food attribute ratings were decodable from the EEG signals during food viewing. A “moving window” approach was implemented to divide each epoch of the EEG data into 10ms segments, which served as analysis time windows. These were moved through the data in steps to derive a fine-grained time-course of decoding performance. The regression model was trained to predict the continuous ratings (ranging from 0-100) for a given food attribute from the spatial brain activity patterns in each time window following the food image onset. A linear SVR model (cost parameter C=0.1, interfacing LIBSVM^74^) was trained on 90% of the data, then tested on the held-out 10% of the data for each time window and for each participant. We repeated this process independently using a 10-fold cross-validation procedure so that all data sets had been used as test data once while training on all other datasets. We repeated this cross-validation process 10 times using newly selected random data to derive a conservative estimate of decoding performance for the time window, calculated as the correlation coefficient between the actual and predicted continuous ratings for the food images, averaged across the 10 × 10 iterations and Fisher-Z transformed. Results for each time window were obtained independently of any other time windows. We used cluster-based permutation tests as described above to correct for multiple tests.

Continuous healthiness and tastiness ratings were decodable from the EEG data (Figure 2E, 2F) from around 150 ms after food image onset, and remained significant until the end of the 1 s analysis window. These findings suggest that there are patterns of activity in the EEG signals that contain fine-grained information about healthiness and tastiness of foods from around 150 ms onwards. Together with the ERP analyses, they show that the FOODEEG dataset is suitable for tracking the first 1000 milliseconds of neural responses to visual food cues.

## Code Availability

All custom code used to produce the technical validation analyses and figures in the manuscript are available from the Open Science Framework (https://doi.org/10.17605/OSF.IO/Y9PMF).

